# Molecular Drivers of RNA Phase Separation

**DOI:** 10.1101/2025.01.20.633842

**Authors:** V Ramachandran, DA Potoyan

## Abstract

RNA molecules are essential in orchestrating the assembly of biomolecular condensates and membraneless compartments in cells. Many condensates form via the association of RNA with proteins containing specific RNA binding motifs. However, recent reports indicate that low-complexity RNA sequences can self-assemble into condensate phases without protein assistance. Divalent cations significantly influence the thermodynamics and dynamics of RNA condensates, which exhibit base-specific lower-critical solution temperatures (LCST). The precise molecular origins of these temperatures remain elusive. In this study, we employ atomistic molecular simulations to elucidate the molecular driving forces governing the temperature-dependent phase behavior of RNA, providing new insights into the origins of LCST. Using RNA tetranucleotides and their chemically modified analogs, we map RNA condensates’ equilibrium thermodynamic profiles and structural ensembles across various temperatures and ionic conditions. Our findings reveal that magnesium ions promote LCST behavior by inducing local order-disorder transitions within RNA structures. Consistent with experimental observations, we demonstrate that the thermal stability of RNA condensates follows the Poly(G) > Poly(A) > Poly(C) > Poly(U) order shaped by the interplay of base-stacking and hydrogen bonding interactions. Furthermore, our simulations show that ionic conditions and post-translational modifications can fine-tune RNA self-assembly and modulate condensate physical properties.

**Author Summary:** RNA molecules are essential for organizing membraneless compartments that play critical roles in cellular processes. While many of these condensates form through interactions between RNA and proteins, recent studies have shown that certain RNA sequences can self-assemble into condensates without protein assistance. This ability is influenced by the sequence composition of RNA and the presence of ions like magnesium. Using detailed molecular simulations we carried out systematic study to reveal how temperature and ionic conditions affect RNA condensation. We discovered that magnesium ions play a key role in driving RNA molecules to condense at lower temperatures by promoting structural changes within the RNA. Our findings also revealed that the stability of RNA condensates varies depending on the RNA sequence, with guanine-rich sequences being the most stable. Additionally, we demonstrated how chemical modifications and ionic conditions can fine-tune the properties of RNA condensates. This study provides new insights into how RNA forms condensates and highlights potential strategies to control their behavior, which could have implications for understanding cellular organization and developing new therapies.

## Introduction

A large fraction of the genome is transcribed into non-coding RNAs (ncRNAs which perform crucial regulatory roles, including transcription, nuclear organization, and catalytic activities (1–3). Over the past decade, numerous experiments have revealed that RNA molecules are dynamically partitioned inside biomolecular condensates or membrane-bound organelles(4–7). Many condensates tend to form via the association of RNA with proteins with specific RNA binding motifs (8–10). However, recent reports have also demonstrated the ability of RNAs to engage in multivalent base-pairing and phase-separation in the absence of proteins. Pure RNA condensates have been observed in CAG, CUG, and other low-complexity repeat sequences(11–14).

Additionally, RNA kissing-loop interactions have been shown to drive the assembly of Oskar ribonucleoprotein (RNP) granules in the female Drosophila germline. Mutations in these loops disrupt granule formation, underscoring the critical role of RNA-RNA interactions in this process (15, 16). The driving forces for RNA phase separation can be broken down into enthalpic contributions from base-pairing and stacking interactions and entropic contributions from collective reorganization of hydration shells and ions accompanying heat or osmotic stress(17–20). The complex interplay of enthalpic and entropic often gives rise to the presence of both Upper Critical Solution Temperature (UCST) and Lower Critical Solution Temperature (LCST), which have been observed in RNA condensates (21–24).

In particular, homopolymeric RNAs such as Poly(G), Poly(A), Poly(C), and Poly(U) have shown base-dependent phase diagrams with different LCSTs. Experiments have found a thermal stability hierarchy among bases, following Poly(G) > Poly(A) > Poly(C) > Poly(U) order (25, 26). The dependence of biomolecular phase separation on ions varies across different systems (27). Divalent ions’ role appears crucial for forming RNA condensates (11, 28, 29). Remarkably, even NaCl alone can induce the phase separation of polyphosphate without the need for divalent ions or crowding agents (30). Similarly, monovalent and divalent ions influence RNA phase separation depending on the nucleotide bases. However, a comprehensive microscopic understanding of how these ions induce phase separation with LCST in RNA condensates has yet to be discovered.

In this work, we employ explicit solvent molecular simulations to unravel the microscopic interactions governing the temperature-dependent phase behavior of RNA condensates and the molecular basis of base-specific LCST phenomena. We systematically analyze homopolymeric RNA tetranucleotides and their chemically modified analogs to map the phase equilibrium landscape across various temperatures and ionic conditions. Our results reveal that magnesium ions promote LCST behavior by driving local order-disorder transitions, facilitating condensate formation. Moreover, our simulations expose a clear thermal stability hierarchy among bases, following Poly(G) > Poly(A) > Poly(C) > Poly(U) order. This ordering highlights the distinct roles of base-specific hydrogen bonding, stacking interactions, and ionic effects in modulating RNA phase separation. Our findings suggest that ionic conditions and chemical modifications can fine-tune RNA self-assembly and alter its phase transitions’ qualitative and quantitative nature. These insights provide a conceptual framework for understanding how cellular factors regulate RNA phase behavior, shedding light on the mechanisms underlying RNA-driven processes in normal physiology and disease contexts.

## Methods

We first generated structures of single-stranded poly G, poly A, poly C, and poly U tetranucleotide chains using the X3DNA library (31, 32). For the m6A4 system, a methyl group was added to the adenine bases using PyMOL. Fifty tetranucleotide chains were then tightly packed within a cubic box of 4.5 nm. This cubic box was subsequently extended into a slab with 4.5 × 4.5 × 25 nm (approximately ∼ 160 mg/ml density). Depending on the specific system, ions (Na+, Cl−, and Mg^2^+) and water molecules were added to fill the box. Using the conjugate-gradient algorithm, the solvated box containing RNA, ions, and water was subjected to energy minimization. The Amber RNA force field parameters (parmBSC0χOL3)(33–35) and the OPC water model were employed (36). For the Mg2+ ions, we used optimized parameters specifically designed for the OPC water model (37), known as microMg, which exhibits a microsecond water exchange rate consistent with experimental observations. The Na+ and Cl− parameters were taken from Young and Cheatham (38). Following energy minimization, we have performed a 4 ns NVT simulation followed by a 4 ns NPT simulation at a temperature of 300 K using the Langevin Middle integrator with a time step of 2 fs (39).

A Monte Carlo barostat, as implemented in OpenMM, was used to maintain an average pressure of 1 bar across all systems. Periodic boundary conditions were applied in all three spatial dimensions. The density of systems was chosen to be approximately 160 mg/ml pragmatically, which is well below the typical condensate phase density (∼500 mg/ml) but significantly higher than the solution phase density used in the experiments (∼1.5mg/ml), thereby facilitating the formation of the stable cluster in a reasonable time. Simulations were run until convergence was achieved, which was assessed by bootstrapping on the time series of cluster sizes using the last two microsecond simulation trajectories (Figure SI 1-7). The DBSCAN algorithm was used to extract the clusters in a frame-by-frame analysis to analyze the size. A detailed contact analysis was also performed using Prolif [37] to investigate the linkages between the cluster characteristics and the various interactions (hydrophobic, electrostatic, and water-bridged interactions) responsible for cluster formation and maintenance.

## Results

### On the role of mono- and divalent ions in RNA condensate formation

Given divalent ions’ crucial role in driving RNA condensate formation, we first dissect the role of ions and ionic interactions on RNA phase separation. To this end, we have constructed three different systems of polyU under varying ionic conditions: (i) neutralization with Na^+^ ions, (ii) neutralization with Mg^2+^ ions, and (iii) neutralization with Na^+^ ions combined with an additional 20 Mg^2+^ ions. We have also simulated these systems at different temperatures, quantifying cluster size and density evolution, which reports on the interplay of enthalpic and entropic forces. Systems with only Na^+^ ions at 270K temperatures form more porous clusters with significant water content, indicating a percolation of RNA (Figure 1A & Figure S8). As the temperature increases to 290K, RNA clusters promptly dissolve, suggesting the presence of an upper critical solution temperature (UCST) (Figure 1A). This narrow window of the phase separation indicates that RNA alone cannot sustain phase separation in the presence of only monovalent ions. However, increasing the length of RNA or introducing crowding agents can help RNA form condensates, as has been demonstrated in experiments (40).

**Figure 1:**
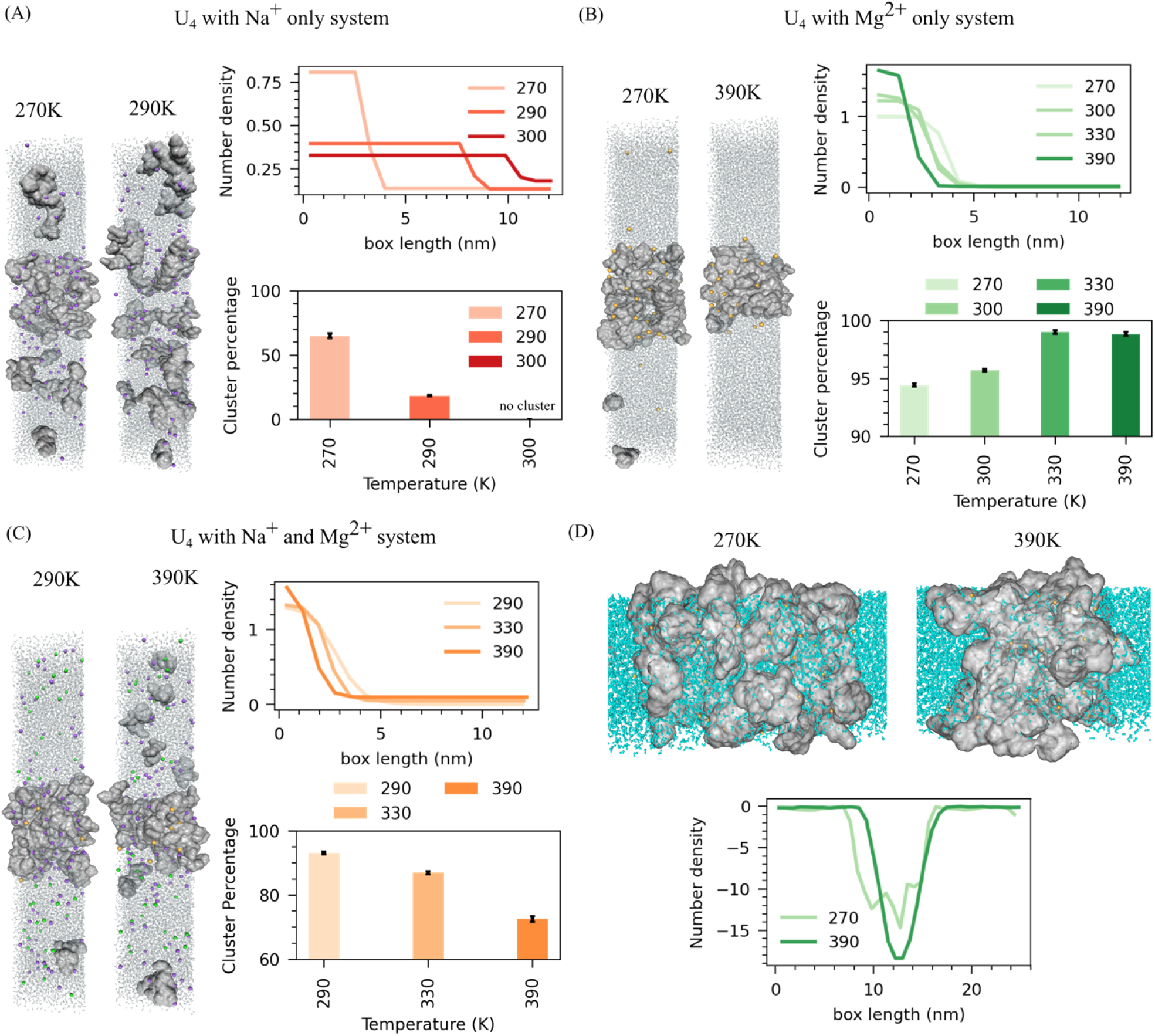
Effects of mono- and divalent ions on RNA condensates. (A) PolyU neutralized with Na^+^ ions: (left) Snapshots from simulations at 270K and 290K. (Right top) Number density profiles of RNA phosphate atoms along the simulation box. (Right bottom) Average RNA Percentage in the cluster. (B) polyU neutralized with only Mg2+ : (left) Snapshot of simulations at different temperatures. (Right top) Number density profiles of RNA phosphate atoms along the box at different temperatures. (Right bottom) Average RNA Percentage in the cluster. (C) polyU neutralized with Na^+^ ions and additional 20Mg^2+^: (Left) Snapshot of simulations at different temperatures. (Right top) Number density profiles of RNA phosphate atoms along the box at different temperatures. (Right bottom) Average RNA Percentage in the cluster. (D) (top) Snapshot of water within clusters at different temperatures and (Bottom) Water number density profiles across the simulation box for polyU neutralized with only Mg2+ ions.

In divalent ion Mg^2+^-only systems, we have chosen a wider range of temperatures (270K, 300K, 330K, and 390K) to conduct simulations because of the larger coexistence window. Unlike monovalent systems, cluster size and density consistently increase with temperature (Figure 1B). At 390K, the RNA clusters become extremely large, dense, and compact, indicating a lower critical solution temperature (LCST) for RNA in the presence of Mg^2+^ (Figure 1B). At the lower temperature of 270K, the cluster remains porous with considerable water content, again highlighting the percolated nature of RNA condensates (Figure 1B and Figure 1D). A similar pattern is also observed when analyzing cluster size; as the temperature increases, one observes the growth of the cluster size (Figure 1B).

Finally, we created a system neutralized with both Na^+^ and 20 Mg^2+^ ions (65mM), which more closely mimics cellular conditions where monovalent ions like Na^+^ and K^+^ are abundant (∼100-150mM), and divalent ions such as Mg2+ are present at lower concentrations. We conducted simulations at T = 290K, 330K, and 390 K. Our results show that increasing temperature leads to a shrinking of cluster size and an increase in density (Figure 1C). Adding a small amount of Mg^2+^ ions promoted RNA phase separation, highlighting the essential role of divalent ions in this process. These findings suggest that even minimal concentrations of Mg^2+^ can significantly impact RNA phase behavior, enhancing its capacity for forming condensates and stabilizing multivalent interactions at lower temperatures.

Overall, in a system containing only Na^+^ ions, a percolated RNA cluster forms at lower temperatures but dissolves as the temperature increases. Conversely, in a system with only Mg^2+^ ions, the cluster persists and becomes larger and denser as the temperature rises (Figure 1 A-C). This behavior can be attributed to two opposing forces: the entropic drive from increased temperature, which tends to dissolve the cluster, and the enthalpic interaction between Mg^2+^ ions and RNA, which promotes cluster formation. A mixed behavior emerges in a mixed system where Na^+^ ions neutralize the charge, and 20 additional Mg^2+^ ions (65mM) are present. As the temperature increases, the cluster density rises while its size decreases (Figure 1 C). These observations suggest that the key factor behind these distinct behaviors is the interaction between Mg^2+^ ions and the phosphate groups of RNA, which intensifies at higher temperatures, driving cluster formation even as the entropic effects push toward dissolution.

Additionally, we find that regardless of ion conditions, at lower temperatures (270K), a significant amount of water becomes trapped within the RNA cluster due to percolation (Figure 1 D & S8-9). As temperature increases, especially in the presence of Mg2+ ions, this water is expelled from the cluster, leading to an entropically favored condensed state. This effect can be quantified by examining the water number density across the simulation box (Figure 1 D & S8-9). Consistent with expectations, water density decreases toward the cluster’s center at all temperatures. Notably, the water density profile displays an irregular well at lower temperatures, whereas at higher temperatures, it sharpens into a distinct single well (Figure 1D).

We next analyzed the ion distribution in all three systems along the box to understand how monovalent and divalent ions drive RNA condensation. The ionic atmosphere analysis shows that both monovalent and divalent ions were densely packed within the RNA clusters (Figure 2 A-B and Figure S10-S12), and notably, Mg^2+^ ion density increased with temperature (Figure 2B). When analyzing the ion interactions within the RNA clusters, specifically Mg^2+^ ions, we observed that outer-sphere Mg^2+^ ions were more prevalent at lower temperatures. However, as the temperature increased, there was a significant rise in the number of inner-sphere Mg^2+^ ions coordinating with one, two, or even three phosphate groups. By 390K, the proportion of Mg^2+^ ions coordinating with phosphate groups had surpassed that of outer-sphere Mg^2+^ ions (Figure 2 C).

**Figure 2:**
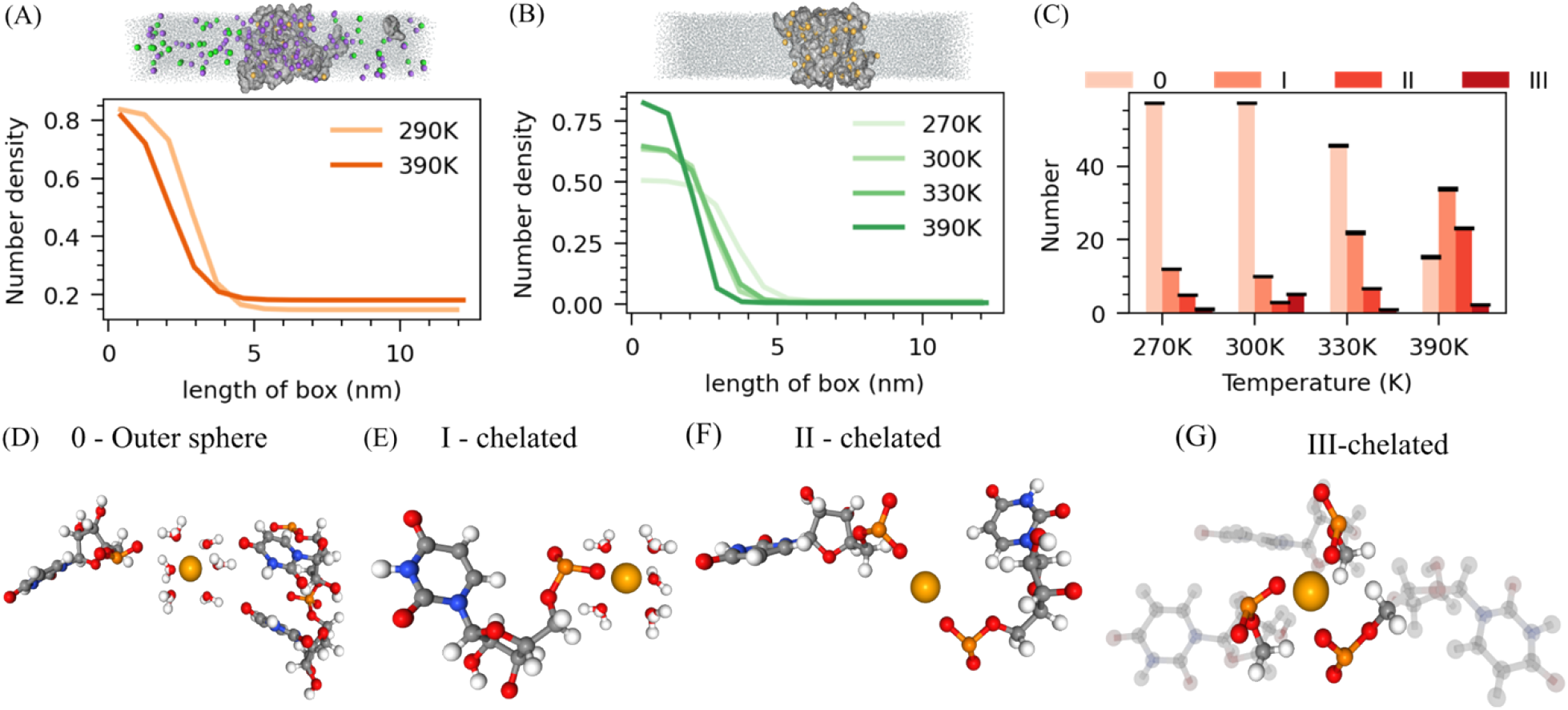
Microscopic-level investigation of the ion atmosphere around RNA. (A) The density of Na^+^ along a box with a snapshot of the simulation of Poly U with Na^+^ and Mg^2+^ ions (B) number density of Mg^2^+ along the box with a simulation snapshot of Poly U with Mg^2+^ only. (C) The average number of outer-sphere, single-chelated, double-chelated, and triple-chelated magnesium is at different temperatures of Poly U with Mg^2+^ only. (D) snapshot of outer-sphere Mg2+, (E) single-chelated Mg^2+^ (F) double-chelated Mg^2+^ and (G) triple-chelated Mg^2+^, interacting with the phosphate backbone of RNA. All snapshots are taken from the Poly U with an Mg^2+^ system only.

This shift in ionic condensation further underscores the role of temperature in driving the coordination of Mg^2+^ with phosphate groups, which, in turn, influences the phase behavior of RNA, particularly its lower critical solution temperature (LCST) in the presence of divalent ions. Increasing temperatures lead to the Mg^2+^ ions driving the water molecules out from the dehydrated shell around phosphate groups, resulting in more compact clusters. This process is particularly evident in the shift from outer-sphere Mg^2+^ ions, where Mg^2+^ is surrounded by water molecules, to inner-sphere Mg^2+^ ions, where Mg^2+^ directly interacts with the phosphate groups, often forming chelation with multiple phosphate groups (Figure 2 C-G). A similar observation was observed in our previous work, where RNA and tetrapeptide of lysine showed reentrant transitions due to desolvation around the phosphate groups of RNA, with increased contacts between phosphate and lysine side chains (23).

### On the role of Base specific interactions on RNA phase-separation

To disentangle the impact of distinct bases chemistry on phase-separation affinities, we conducted phase coexistence simulations for four separate systems, each containing 50 chains (approximately 160 mg/ml) of one type of tetranucleotide: guanine (G4), adenine (A4), cytosine (C4), and uracil (U4). Na^+^ ions were included in each system for charge neutralization, along with 20 additional Mg^2+^ ions (65mM). This ion setup mimics cellular conditions while minimizing phosphate chelation through limited Mg^2+^ ions, enhancing the differentiation in phase separation behavior driven by the bases. These simulations were conducted at two temperatures, 330K and 390K, to examine temperature-dependent effects. Our findings reveal that the size and characteristics of the formed clusters strongly depend on the RNA bases.

At T=330K, Guanine and Adenine form larger clusters, with most residues in the cluster, than Cytosine and Uracil, which form comparable cluster sizes. The cluster size order follows Poly G ∼ Poly A > Poly C ∼ Poly U (Figure 3 C). As temperature rises to 390K, Uracil’s cluster size decreases faster, and cluster size follows: PolyG > Poly A> Poly C > Poly U (Figure 3 C). In all bases, cluster size decreases with temperature rises while cluster density increases (Figure 3C × Figure S13). At both temperatures, 330K and 390K, the condensate state density of all bases is approximately the same. This is because Mg^2+^ ions play a central role in condensate formation through chelation with the phosphate group. When comparing the condensate slab width, it follows Poly G> Poly A > Poly C>Poly U in both temperatures (Figure 3A-B and Figure S13 A-B). Also, the vapor phase density in both temperatures shows that guanine and adenine exhibit lower cluster densities than cytosine and uracil. Therefore, the overall condensation ability is Poly G > Poly A > Poly C > Poly U (Figure S13 A-B).

**Figure 3:**
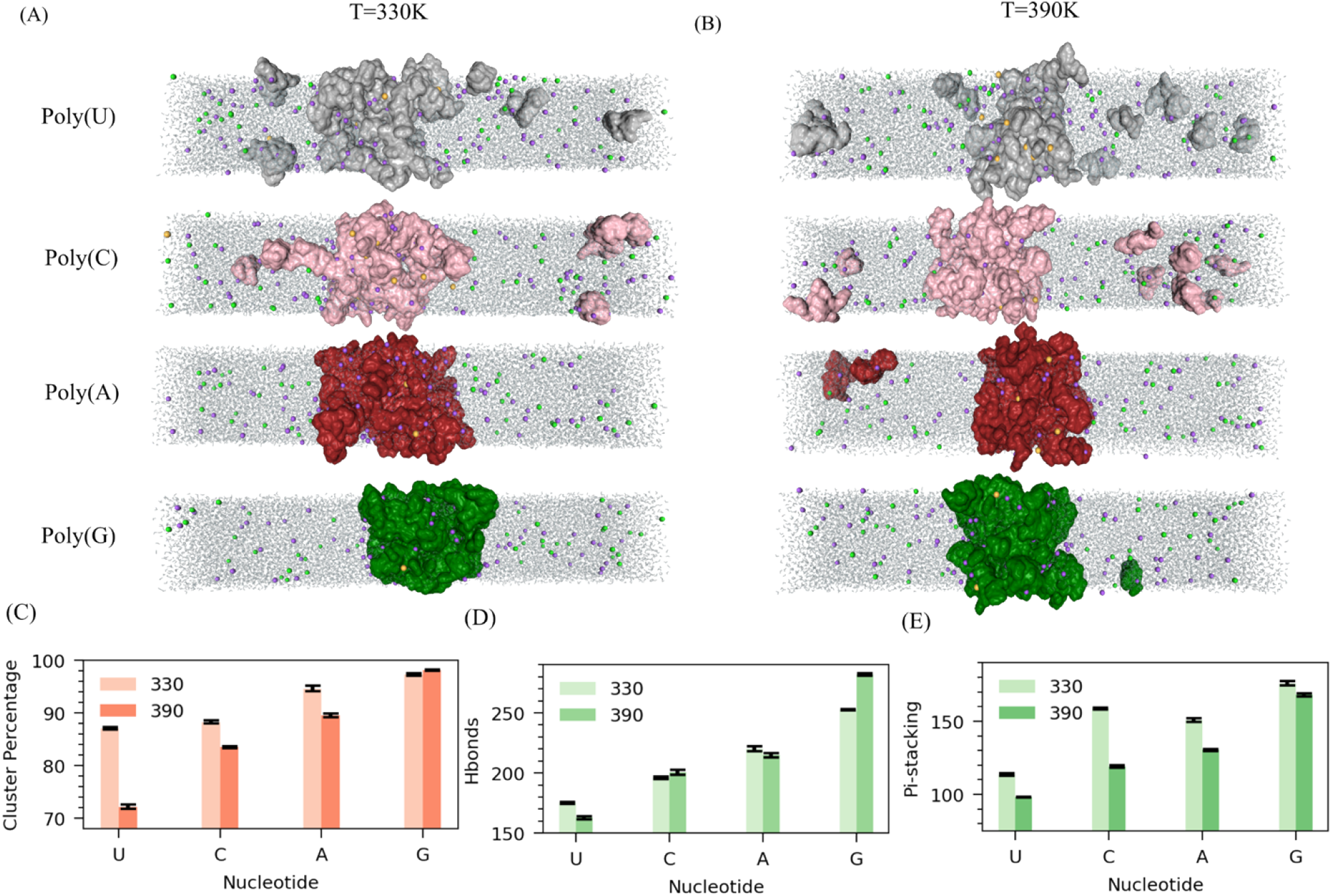
Influence of base chemistry in RNA LLPS. Snapshot from Poly G, Poly A, Poly C, and Poly U slab simulation at two temperatures: (A) 330K and (B) 390K. (C) Average RNA percentage in the cluster comparison among Poly G, Poly A, Poly C, and Poly U at two temperatures 330K and 390 K. Microscopic interactions: (D) Hbonds and (E) Pi-stacking comparison among Poly G, Poly A, Poly C, and Poly C at two temperatures. All systems are neutralized here with Na^+^ ions and 20 additional Mg^2+^ ions.

We calculated microscopic interactions, including hydrogen bonding, pi-stacking, and base pairing, to delve deeper into the base-dependent phase separations. At both temperatures, the trend for hydrogen bonds strictly follows Poly G > Poly A > Poly C > Poly U (Figure 3D). Similarly, pi-stacking follows the same trend. At lower temperatures T=330K, G4, A4, and C4 form a larger number pi - stacking interactions than Uracil (Figure 3E). Guanine forms the most extensive hydrogen bonds and pi-stacking interactions. In terms of base pairing, guanine again forms the highest number of pairs, followed by adenine and uracil, with cytosine forming the fewest pairs (G4 > A4 > U4 > C4) (Figure S13 C).

Notably, guanine and adenine exhibit a higher propensity for base pairing than cytosine and uracil. Overall, the interaction strength across all microscopic parameters follows the trend: PolyG > PolyA > PolyC > PolyU, aligns well with the observed condensation tendencies of these polynucleotides, where guanine-rich sequences display the strongest phase separation behavior, followed by adenine, cytosine, and uracil.

We divided each nucleotide into its phosphate, sugar, and base groups to further investigate the specific interactions driving RNA phase separation. We calculated the radial distribution function (RDF) for each pair. Our analysis revealed that guanine exhibits the strongest phosphate-base interactions, followed by cytosine, owing to the presence of the NH_2_ group in the base. The phosphate-base interaction strength follows the order: G_4_ > C_4_ > A_4_ > U_4_ in both temperatures 330K and 390K (Figure 4A & Figure S14). Adenine prefers to form strong base-base interactions through the NH2 group in its base, as reflected in its RDF, with the base-base interaction strength following the order of G4 > A4 > C4 > U4 (Figure 4B & 4E). Guanine’s ability to form strong phosphate-base and base-base interactions explains its enhanced phase separation potential, whereas uracil, lacking precise specific interactions, exhibits a weaker tendency to phase separate.

**Figure 4:**
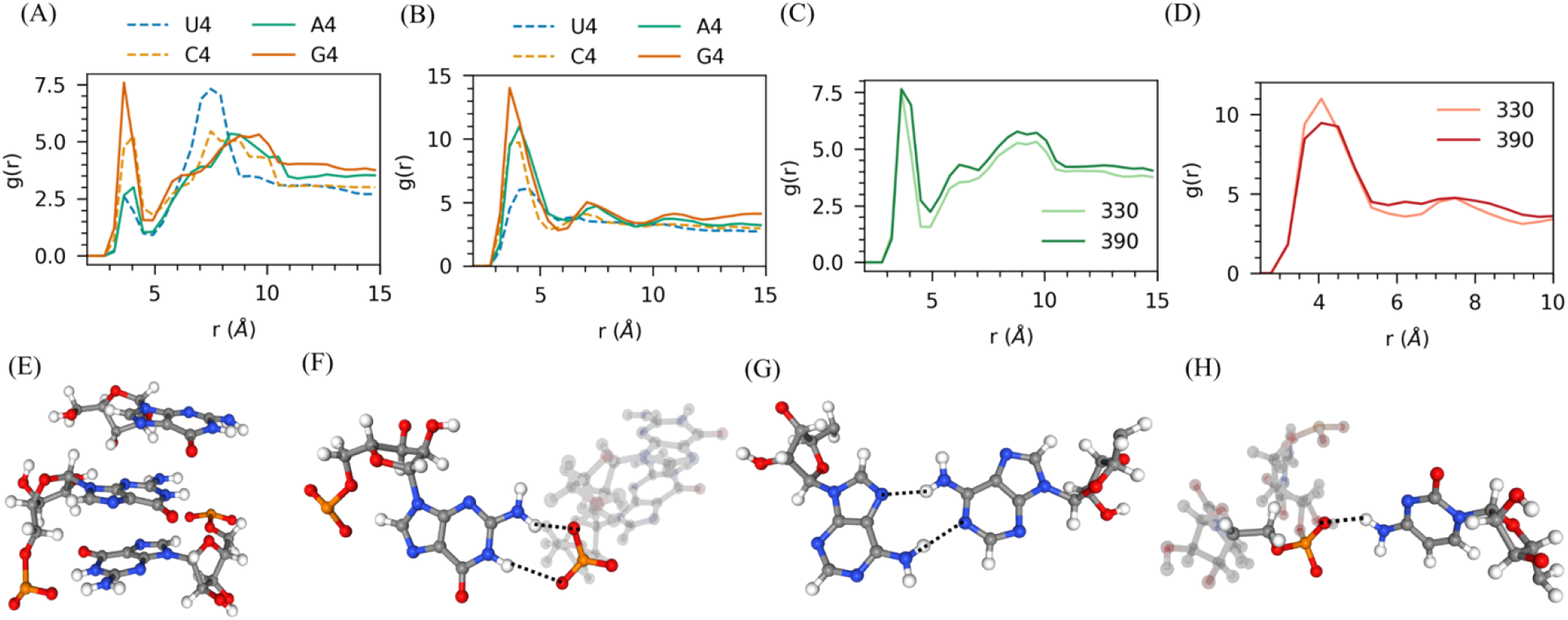
Investigation of specific interactions driving homopolymer RNA phase separation using RDF. RDF of (A) phosphate-base and (B) base-base. (C) phosphate-base rdf at two temperatures of guanine and (D) base-base rdf at two temperatures of adenine. (E) Snapshot of pi-stacking interactions in Guanine. (F) Hbond pairs between phosphate and base in guanine. (G) Base pair in (hydrogen bonds between pairs between bases) in Adenine. (F) hydrogen bond between phosphate and base in cytosine. Here, all systems are neutralized with Na^+^ ions along with 20 additional Mg^2+^ ions.

When examining the temperature dependence, in all four bases, phosphate-base interactions increase with rising temperature, while base-base interactions decrease (Figure 4F & 4G & Figure S15). This observation underscores the role of phosphate in driving the Lower Critical Solution Temperature (LCST) behavior. Lastly, we analyzed the interactions of Na^+^ ions around the phosphate and base groups of RNA. Across all four bases, the Na^+^ distribution around the phosphate groups increased with rising temperature. Around the RNA base, Na^+^ distribution also increased with temperature in all cases except adenine (Figure S16 and S17). Interestingly, the interaction of Na^+^ with the adenine base decreases as the temperature rises (Figure S16 and S17). At both temperatures, the distribution of Na^+^ around both the phosphate and base groups followed the trend: Poly G > Poly C ∼ Poly U > Poly A. This suggests that adenine interacts less with ions than other nucleotides due to its base’s absence of a C=O group.

### Effects of Adenine Methylation on RNA LLPS

The cellular environment is a highly dynamic and chemically active space where proteins and RNA undergo various post-translational and post-transcriptional modifications, such as phosphorylation, methylation, and acetylation (41, 42). These chemical modifications shape biomolecules’ behavior, dynamics, and phase separation properties. In this study, we have extensively explored the effects of ions and base chemistry on RNA’s ability to undergo condensation and phase separation. Equally important, however, is understanding how chemical modifications on RNA impact these processes. RNA can undergo several types of methylation, with N6-methyladenine (m6A) being one of the most common modifications involved in various cellular processes (Figure 5A-B) (43, 44).

**Figure 5:**
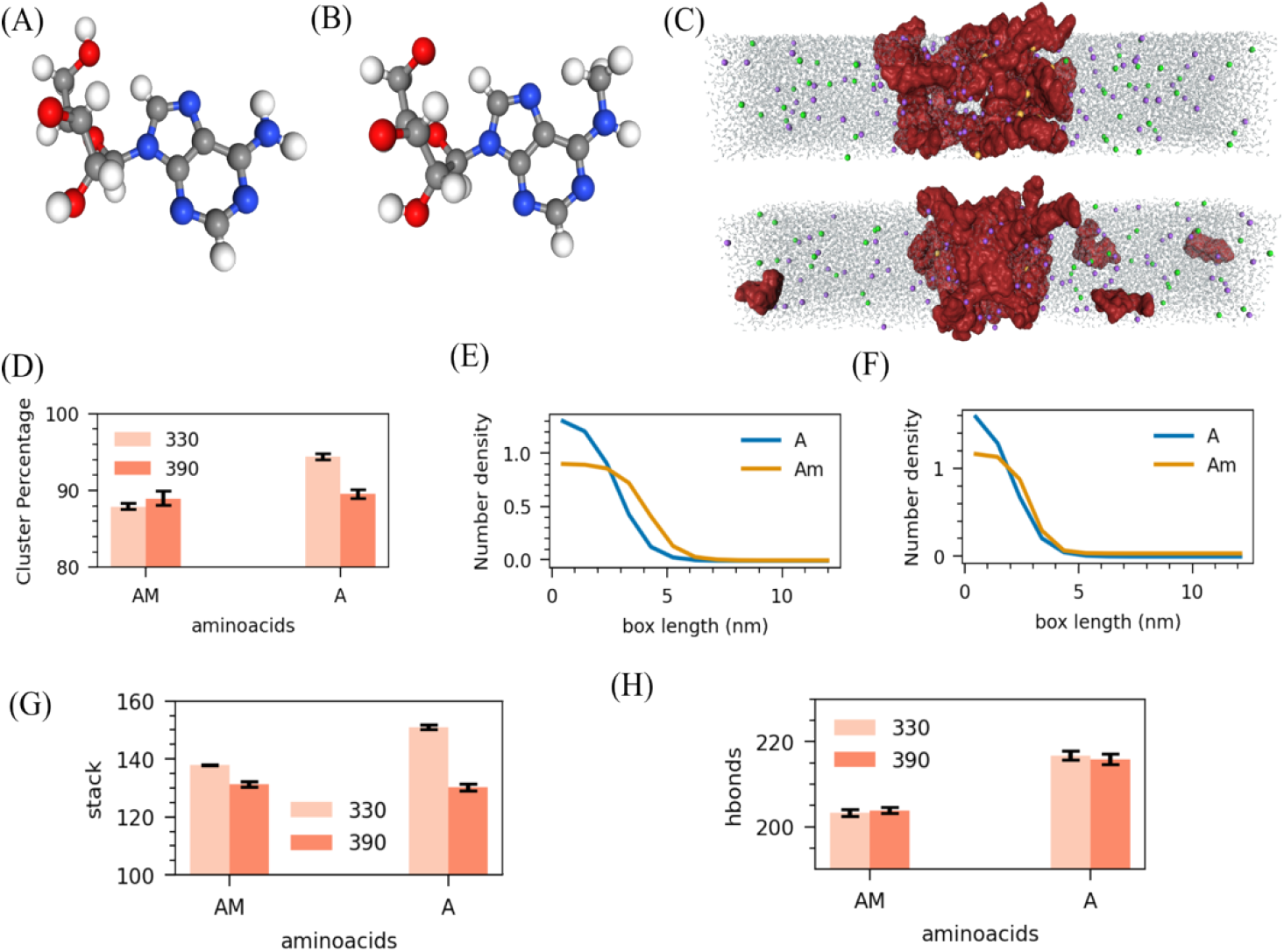
Effects of N6-methylation on RNA interactions and its role in liquid-liquid phase separation (LLPS). (A) Adenine base structure (B) N6-methyladenine base (C) Snapshots from simulations at 330K (top) and 390K (bottom). Comparison of (D) Average RNA percentage in the cluster and Number density of phosphate along a box at (E) 330K and (F) 390K of m6A4 and A4. Comparison of microscopic interactions: (G) total number of pi-stacking and (H) total number of hydrogen bonds. Here all systems are neutralised with Na^+^ ions along with 20 additional Mg^2+^ ions.

To investigate the effects of N6-methylation on RNA interactions and its role in liquid-liquid phase separation (LLPS), we performed slab simulations containing 50 chains of m6A-modified RNA tetramers (m6A4, approximately 160 mg/ml) in the presence of Na^+^ ions for charge neutralization with 20 additional Mg^2+^ ions (65mM) at two different temperatures: 330K and 390K. To assess the impact of N6-methylation on RNA condensation, we calculated the cluster density and size and compared it with A4.

We find that m6A4 forms smaller clusters at both temperatures than A4 (Figure 5D). Additionally, across both temperature conditions, the clusters formed by m6A4 are less dense than those formed by A4 (Figure 5E-F).

These findings suggest that the m6A modification suppresses phase separation, reducing the overall condensation ability of RNA. We further examined microscopic interactions: hydrogen bonding, pi-stacking, and base pairing. Our results reveal that the m6A modification reduces overall hydrogen bonding more than A4 at 330K and 390K. Similarly, m6A exhibits lesser pi-stacking interactions and fewer hydrogen bonds than A4. (Figure 5G-H and Figure S18). It explains that substituting NH2 with NHCH3 in m6A hinders the hydrogen bonding and Pi-stacking interaction ability, suppressing RNA condensates’ stability.

## Discussion

RNA-based condensates have been shown to play key roles in transcriptional regulation, catalytic activities, stress response, and sensing functions [35–38]. Much of the recent focus has been on protein components sequestering RNAs into condensates. However, recent experiments have discovered an innate ability of RNA to form condensate phases without much protein assistance. However, a detailed understanding of how homotypic RNA interactions contribute to the formation and properties of such condensates remains to be fully explored [34]. In this study, we employ atomistic molecular simulations to elucidate the molecular driving forces governing the temperature-dependent phase behavior of RNA, providing new insights into the origins of LCST. Using RNA tetranucleotides and their chemically modified analogs, we map RNA condensates’ equilibrium thermodynamic profiles and structural ensembles across various temperatures and ionic conditions.

Four distinct RNA tetranucleotides U4, G4, C4, and U4 were simulated under varying temperatures (270K to 390K, depending on the system) and salt conditions to explore how ions, base chemistry, and chemical modifications affect RNA phase behavior. We first focused on the ionic environment’s impact on RNA LLPS, specifically using poly uracil (Poly U) in three ionic conditions: (i) neutralized with Na^+^ ions, (ii) neutralized with Mg^2+^ ions, and (iii) neutralized with Na^+^ ions plus an additional 20 Mg^2+^ ions (65mM). Results showed that, in the presence of Na^+^ alone, RNA formed percolated clusters at lower temperatures, which dissolved as temperatures increased. Conversely, in systems with Mg^2+^ alone, clusters grew larger and denser with rising temperature, indicating Lower Critical Solution Temperature (LCST) behavior. Interestingly, RNA clusters retained considerable water at lower temperatures across all ionic conditions, with water gradually expelled as the temperature rose, suggesting that entropically driven dehydration promotes condensation. We used an RNA density of approximately 160 mg/mL—higher than typical experimental values—to ensure stable cluster formation, leading to percolated structures at lower temperatures.

Our subsequent analysis of the molecular interaction of RNA ions shows that outer-sphere Mg^2+^ ions were more prevalent at lower temperatures, loosely interacting with the RNA. However, as the temperature increased, there was a notable rise in the number of inner-sphere Mg^2+^ ions, which began to coordinate directly with one, two, or even three phosphate groups, suggesting that the increase in entropy due to temperature favors Mg^2+^ chelation with the phosphate groups. Higher temperatures are required for the phosphate to shed its hydration shell, allowing it to replace the first layer of water molecules and chelation of Mg^2+^, highlighting how temperature drives the interaction network within the condensate through Mg^2+^ ions. A similar mechanism is observed in our previous Poly U-K4 reentrant transition study, where desolvation of phosphate groups at higher temperatures led to increased phosphate-lysine interactions. Thus we conclude that water expulsion and Mg^2+^ chelation are key energetic and entropic drivers for RNA condensate formation.

Next, we have explored how the chemical nature of the nucleobases affects RNA LLPS by simulating guanine (G4), adenine (A4), cytosine (C4), and uracil (U4) systems in the presence of Na^+^ for neutralization and 20 additional Mg^2+^ ions (65mM), at two temperatures (330K and 390K). Our findings align with experiments(25), showing that the overall condensation ability follows the trend: PolyG > PolyA > PolyC > PolyU. By carrying out a detailed analysis of patterns of hydrogen bonding, π-stacking, and base pairing in RNA condensates we found that the strength of all these interactions follows the same trend hierarchy PolyG > PolyA > PolyC > PolyU. Guanine exhibits the strongest phosphate-base interactions, likely due to the NH2 group on its base. Cytosine follows closely, while adenine favors strong base-base interactions through its NH2 group. These insights align with our previous computational studies on 20-polynucleotide sequence repeats(26), which showed that strong Pi-stacking interactions combined with hydrogen bonds between bases and phosphates can cause well-organized stems or hairpin loop structures in PolyG. In PolyA, base pairing dominates, driving the formation of stem and pseudoknot structures, while in PolyC, hydrogen bonding between the NH2 groups of the base and phosphate is prominent. In contrast, PolyU shows minimal interactions, resulting in a highly mobile structure. Interestingly, we observed that phosphate-base interactions increase with rising temperature. Conversely, base-base interactions decrease, highlighting the pivotal role of phosphate in driving LCST behavior. Additionally, adenine interacts less with ions than other nucleotides due to its base’s absence of a C=O group.

Finally, we have examined the effect of post-translational modifications on RNA phase separation, focusing on N6-methyladenine (m6A), a common RNA modification. Simulations of m6A-modified RNA tetramers (m6A4) at 330K and 390K showed that substituting the NH2 group with NHCH3 in m6A reduced hydrogen bonding potential, weakening LLPS. It suggests that m6A modification suppresses phase separation compared to unmodified adenine. Interestingly, experimental studies show that m6A modifications enhance RNA-protein interactions [35]. For example, m6A-binding proteins—YTHDF1, YTHDF2, and YTHDF3—undergo LLPS in vitro and in cells, with phase separation significantly amplified by mRNAs containing multiple m6A residues [36]. Through this study, we aimed to provide a comprehensive understanding of the factors that influence RNA-RNA interactions in condensate design. These insights are expected to enhance our understanding of the mechanisms underlying the formation of various cellular membraneless condensates.

## Data Availability

The data underlying this article are available in the article and in its online supplementary material.

## Supporting Information

Figure S1-S18

## Funding

Authors acknowledge support from the National Institutes of Health with grant no R35GM138243.

## Conflict of interest statement

None declared.

## Notes

### Competing Interest Statement

The authors have declared no competing interest.

